# Structure and mechanism of a primate ferroportin

**DOI:** 10.1101/2020.03.04.975748

**Authors:** Zhenning Ren, Shuai Gao, Jiemin Shen, Lie Wang, Zhichun Xu, Ye Yu, Preetham Bachina, Hanzhi Zhang, Arthur Laganowsky, Nieng Yan, Ming Zhou, Yaping Pan

**Affiliations:** Verna and Marrs McLean Department of Biochemistry and Molecular Biology, Baylor College of Medicine, Houston, TX 77030, USA; Department of Molecular Biology, Princeton University, Princeton, NJ 08544, USA; Department of Chemistry, Texas A & M University, College Station, TX 77843, USA

**Author notes:** These authors contributed equally to the work (Zhenning Ren, Shuai Gao, Jiemin Shen, Yaping Pan). Correspondence to M. Zhou and Y. Pan.

## Abstract

Ferroportin is the only cellular iron exporter in human and essential for iron homoeostasis. Mutations in ferroportin are associated with hemochromatosis or ferroportin diseases characterized by a paradoxical combination of anemia and abnormal accumulation of iron in cells. Ferroportin is also the target of hepcidin, which is a hormone that downregulates ferroportin activity. However, due to a lack of three-dimensional structures, the mechanism of iron transport by ferroportin and its regulation by hepcidin remains unclear. Here we present the structure of a ferroportin from the primate Philippine tarsier (TsFpn) at 3.0 Å resolution determined by cryo-electron microscopy. TsFpn has a structural fold common to major facilitator superfamily of transporters and the current structure is in an outward-open conformation. The structure identifies two potential ion binding sites with each site coordinated by two residues. Functional studies demonstrate that TsFpn is a H^+^/Fe^2+^ antiporter and that transport of one Fe^2+^ is coupled to the transport of two H^+^ in the opposite direction such that the transport cycle is electroneutral. Further studies show that the two ion binding sites affect transport of H^+^ and Fe^2+^ differently. The structure also provides mechanistic interpretation for mutations that cause ferroportin diseases.

## Introduction

In mammals, ferroportin (Fpn) exports cellular iron and is highly expressed in enterocytes, hepatocytes and macrophages to distribute iron absorbed from food or recovered from digestion of senescent red blood cells^1^. Fpn also mediates iron transport across the placenta and thus is required for the normal development of embryos^2^. Fpn activity is regulated by hepcidin, a peptide hormone, which reduces Fpn activity by a combination of inhibiting the transport activity and promoting endocytosis of Fpn^3,4^. More than sixty Fpn mutations have been identified in human to cause ferroportin diseases^5,6^ that are characterized by accumulation of iron inside of macrophages, highlighting its important physiological role in iron homeostasis. Fpn and its modulation by hepcidin has been the focus of targeted therapeutics for treating ferroportin diseases^7–9^.

Fpn belongs to the solute carrier family 40 (SLC40A1)^10–12^ and is a member of the major facilitator superfamily (MFS) of secondary transporters which includes the glucose transporter (GLUT1-5)^13–15^, peptide transporter (PEPT1, SLC15A1)^16–18^, and equilibrative nucleoside transporter^19^. Transporters of the MFS family share a common structural fold that has two homologous halves forming a clam-shell like architecture. A single substrate binding site is commonly located to the center of the clam-shell, and substrate translocation is achieved by rock-switch type motions of the two halves of the clam-shell so that the substrate binding site is alternatively exposed to either side of the membrane^20^. Structures of a bacterial homolog of Fpn (*Bdellovibrio bacteriovorous*; BbFpn) were reported recently^21,22^, which enhances our understanding of Fpn. However, BbFpn has ~20% sequence identity and 49% similarity to that of human Fpn and may not depict an accurate representation of the mechanism of iron recognition and transport in human Fpn. We expressed and purified an Fpn from Philippine tarsier (*Tarcius syrichta* or *Carlito syrichta*; TsFpn), which is 92% identical and 98% similar to human Fpn, and we characterized its function and determined its structure.

### In vitro functional studies of TsFpn

TsFpn is expressed and purified from insect cells and elutes as a single peak on a size-exclusion chromatography column. The elution volume is consistent with TsFpn being a monomer (Fig. 1a and Methods). TsFpn has three predicted N-linked glycosylation sites^23^ and migrates as a diffused band on an SDS-PAGE, and addition of glycosidases PNGase F and EndoH helps focus the protein band (Fig. 1a), confirming that TsFpn is glycosylated.

**Figure 1.**
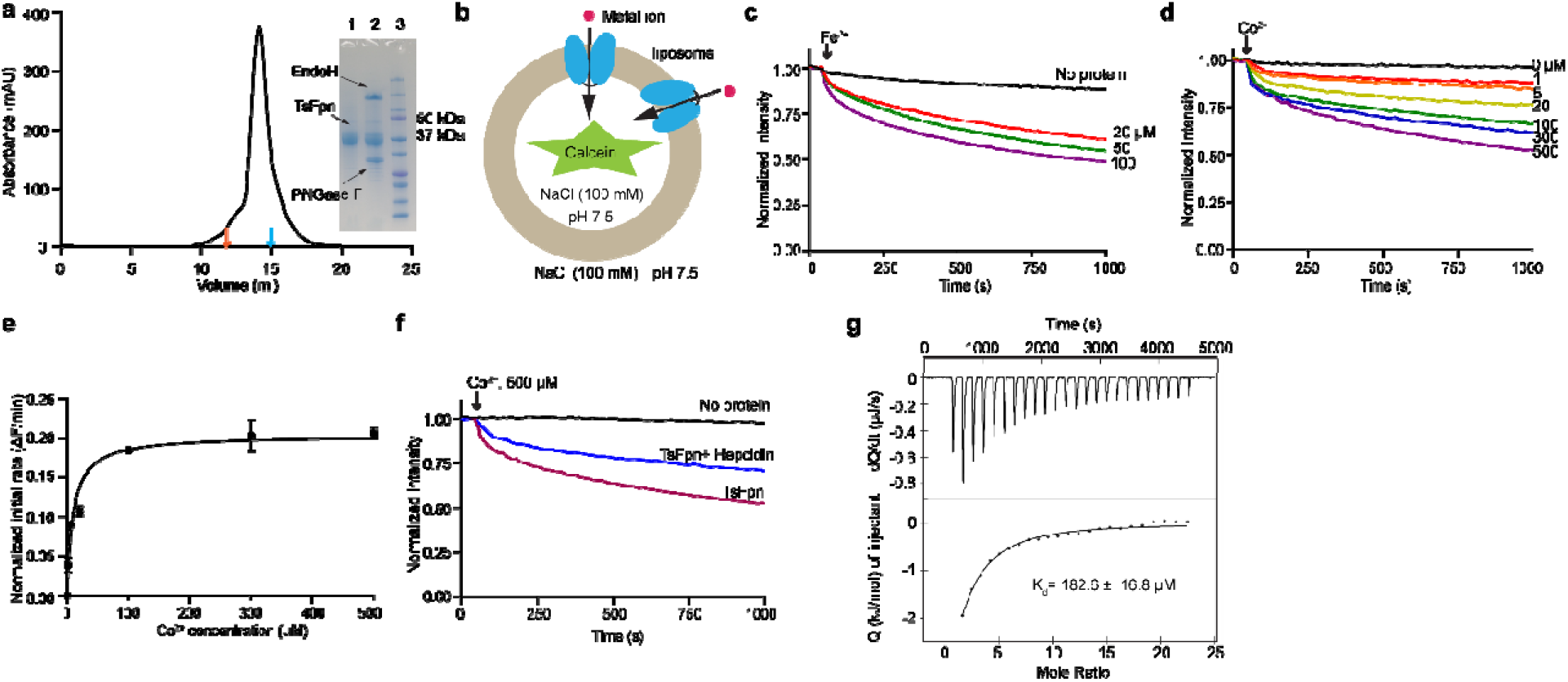
Function of purified TsFpn. **a.** Size-exclusion chromatography profile of TsFpn in detergent DDM. Elution volumes of membrane proteins of known molecular weight, bcMalT (100 kDa, orange)^48^ and mouse SCD1 (41 kDa, blue)^32^ are marked by arrows. Inset: SDS-PAGE of purified TsFpn (Lane 1), purified TsFpn with EndoH glycosidase (Lane 2), and standard molecular weight marker (Lane 3). Molecular weights of two the markers are labeled on the right. **b.** Schematic view of a TsFpn containing proteoliposome with calcein dye enclosed. Influx of metal ions (Fe^2+^ or Co^2+^, red spheres) quenches the fluorescence. Two orientations of TsFpn are shown. **c.** Quench of fluorescence in the presence of various concentrations of external Fe^2+^ at the presence of 1 mM vitamin C for liposomes with no protein (black line) or liposomes with TsFpn. **d.** Quench of fluorescence in the presence of different concentrations of external Co^2+^. **e.** Initial rate of fluorescence quench versus concentration of Co^2+^. The solid line is the data fit to a Michaelis Menten equation. **f.** Quench of fluorescence in the presence of 500 μM Co^2+^ for liposomes with no protein (black line), proteoliposomes with TsFpn (red line), or proteoliposomes with TsFpn and with 1 μM hepcidin added to the external side of the liposomes (blue line). **g.** ITC measurement of Co^2+^ binding to TsFpn. Each spike is rate of heat release (upper panel) and each point is the integration of the spike (lower panel). The solid line in the lower panel is fit of the data to a single binding site. Each data point in **e** is the average of 3 or more measurements, and the error bars are standard error of the mean (s.e.m.).

We reconstituted the purified TsFpn protein into liposomes and measured transport of Fe^2+^ using a flux assay (Fig. 1b and Methods). Liposomes reconstituted with TsFpn show significant Fe^2+^ uptake while liposomes without the protein do not have significant change of fluorescence over time (Fig. 1c). Since Fe^2+^ is easily oxidized under aerobic conditions, a reducing reagent (1 mM vitamin C) was added to the external solution to stabilize ferrous. Although ferrous transport can be observed, the addition of reducing reagents affects free Fe^2+^ concentrations thereby complicating the measurement. Thus, Co^2+^ was used to further characterize the transport activity of TsFpn. TsFpn mediates Co^2+^ uptake and we measured the uptake at different Co^2+^ concentrations. The initial rate of uptake versus ion concentration can be fit with a Michaelis Menten equation with a K_M_ of 9.7 ± 3.26 μM and V_max_ of 0.20 ± 0.03 ΔF/min (Fig.1d, e). TsFpn is sensitive to human hepcidin^3^, and the inhibition reaches ~50% likely due to the random orientation of TsFpn on the liposomes (Fig. 1f). We also measured Co^2+^ binding to the purified TsFpn using isothermal titration calorimetry (ITC) and found that the binding is exothermic with a ΔH of −12.0 ± 0.55 kJ/mol and TΔS of 9.29 ± 0.38 kJ/mol. TsFpn binds to Co^2+^ with a dissociation constant (K_d_) of 182.6 ± 16.8 μM (Fig. 1g).

### Structure of TsFpn

We prepared monoclonal antibodies against TsFpn to facilitate its structural determination^24^ (Methods). TsFpn forms a stable complex with the antigen binding fragment (Fab) of a monoclonal antibody 11F9 as indicated by a shift in the retention time of the elution peak on the size-exclusion column (Extended Data Fig. 1a-b). To assess the effect of 11F9 Fab on TsFpn, we examined Co^2+^ binding and transport in the presence of the Fab. Fab-TsFpn complex has a modestly reduced affinity to Co^2+^ with a K_d_ of 258.2 ± 34.2 μM (Extended Data Fig. 2a). However, addition of the Fab inhibits Co^2+^ uptake, and the inhibition reaches ~50% at 1 μM of Fab (Extended Data Fig. 2b-c). These results suggest that the Fab has a modest effect on ion binding and a pronounced effect on ion transport, likely by hindering conformational changes of TsFpn.

We reconstituted TsFpn-11F9 into nanodisc (Extended Data Fig.1a) and prepared cryo-EM grids in the presence of 10 mM of Co^2+^. The images show recognizable particles of TsFpn-11F9 complex and we were able to obtain a final map at 3.0 Å overall resolution (Fig. 2a, Extended Data Figs. 3 and 4 and Extended Data Table 1). The map shows clear density for all transmembrane helices and resolves most of the side chains (Extended Data Fig. 5) to allow *de novo* building of the TsFpn structure. The final structure model includes residues 17 to 237, 289 to 395 and 453 to 552. The N-terminal 16 residues, two loops between TM6 and 7 and TM9 and 10, and C-terminal 25 residues were not resolved. These regions are predicted to be unstructured (Extended Data Fig. 6). For the Fab fragment, the constant region was not fully resolved and was built as poly alanines while the variable region is well resolved with a local resolution close to 2.9 Å (Extended Data Fig. 3).

**Figure 2.**
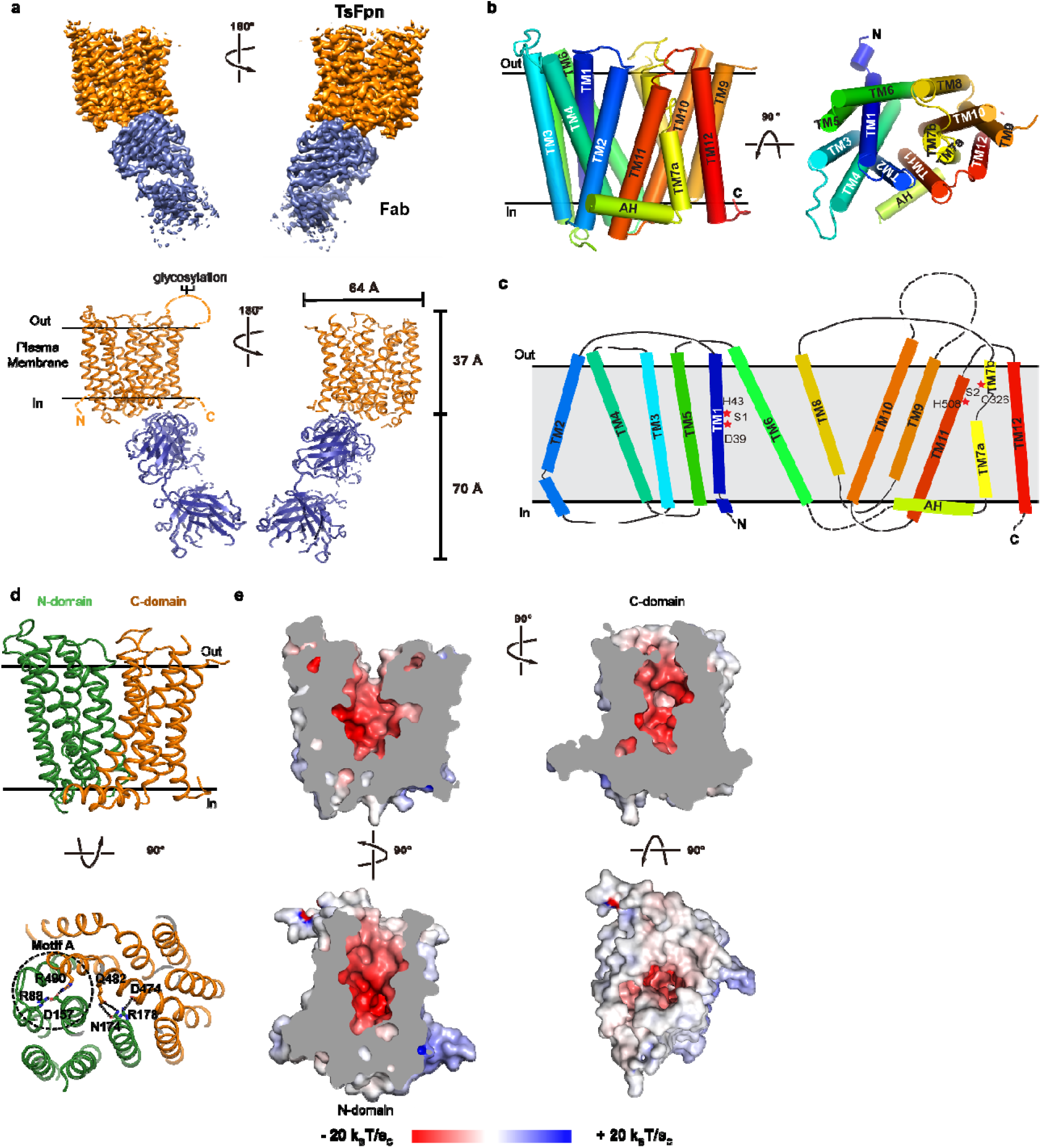
Structure of TsFpn. **a.** Top panel: cryo-EM map of TsFpn (orange) in complex with Fab (blue) in two views. Bottom panel: TsFpn in complex with Fab shown as ribbon representation. **b.** TsFpn structure shown as cylinder representation and in two views. **c.** topology of TsFpn. Regions that are not resolved in the structure are marked as dotted lines. **d.** Top: The N- and C-domains of TsFpn shown in green and orange, respectively. Bottom: TsFpn viewed from the intracellular side. Interacting residues from the N- and C-domains are marked as sticks. **e.** Electrostatic static potential of TsFpn mapped onto the surface representation. The cut-away views show the large cavity formed between the N- and C-domains. Electrostatic static potential is calculated by APBS^49^ in Pymol.

TsFpn adopts a canonical MFS fold^25^. The 12 transmembrane helices are packed into two clearly defined domains. TM1-6 form the N-domain, and TM7-12 the C-domain (Fig. 2b-d). Based on previous studies of human Fpn topology^26^ and the “positive-inside” rule^27^, both the N- and C-termini are located to the cytosolic side. The two domains are connected by a long intracellular loop between TM6 and TM7. Part of the loop is an amphipathic helix that extends horizontally and oriented parallel to the intracellular surface of the membrane (Fig. 2b).

TsFpn structure is in an outward facing conformation. The N- and C-domains make contact at the cytosolic side. Asp157 on TM4 is in close proximity to Arg88 on TM3 and Arg490 on TM11 and could form salt bridges with the arginines (Fig. 2d). These residues are conserved in the MFS family of transporters and are commonly known as the motif-A^25^. Other interactions between the N- and C-domains include Arg178 in the N-domain and Asp474 in the C-domain, and Asn174 in the N-domain and Gln482 in the C-domain. These interacting pairs of residues are conserved in human Fpn (Extended Data Fig. 6) and mutations to Arg88, Asp157, Asn174, Arg178, and Arg490 are known to cause ferroportin diseases^5,28,29^.

### Potential metal ion binding sites

TsFpn structure has a large solvent-accessible central cavity between the N- and C-domains (Fig. 2e). Residues lining the cavity are mostly charged or hydrophilic, and although there are several arginine residues, the electrostatic surface potential of the cavity is highly negative (Fig. 2e). Two strong non-protein densities stand out in the cavity and we assigned the two as potential metal ion binding sites. The first site, S1, is coordinated by Asp39 and His43, and the second site, S2, Cys326 and His508 (Fig. 3a-c). S1 is located in the N-domain while S2 is in the C-domain. The densities at S1 and S2 are comparable to surrounding residues (Fig. 3b-c). Both S1 and S2 are solvent accessible from the extracellular side, and the distance between the two sites are 16.0 Å as measured between the two ions. These two metal ion binding sites are unusual because both the S1 and S2 sites are coordinated by only 2 residues, which is very different from the ion binding sites identified in other transition metal ion transporters of known structures, such as NRAMP^30,31^, VIT1^31^, YiiP^32^, ZIP^33^ and ZneA^34^, all of which have at least four residues coordinating a metal ion binding site. In addition, S2 does not have a charged residue making direct contact with the bound ion, although two negatively charge residues, Asp325 and Asp505 are located close to S2 and could potentially interact with Cys326 and His508, respectively. It is also unusual to have two substrate binding sites because most other members of the MFS family of transporters have a single substrate binding site often coordinated by residues from both the N- and C-domains^13–19^. To further understand how S1 and S2 may participate in ion transport, we examined the binding sites with functional studies.

**Figure 3.**
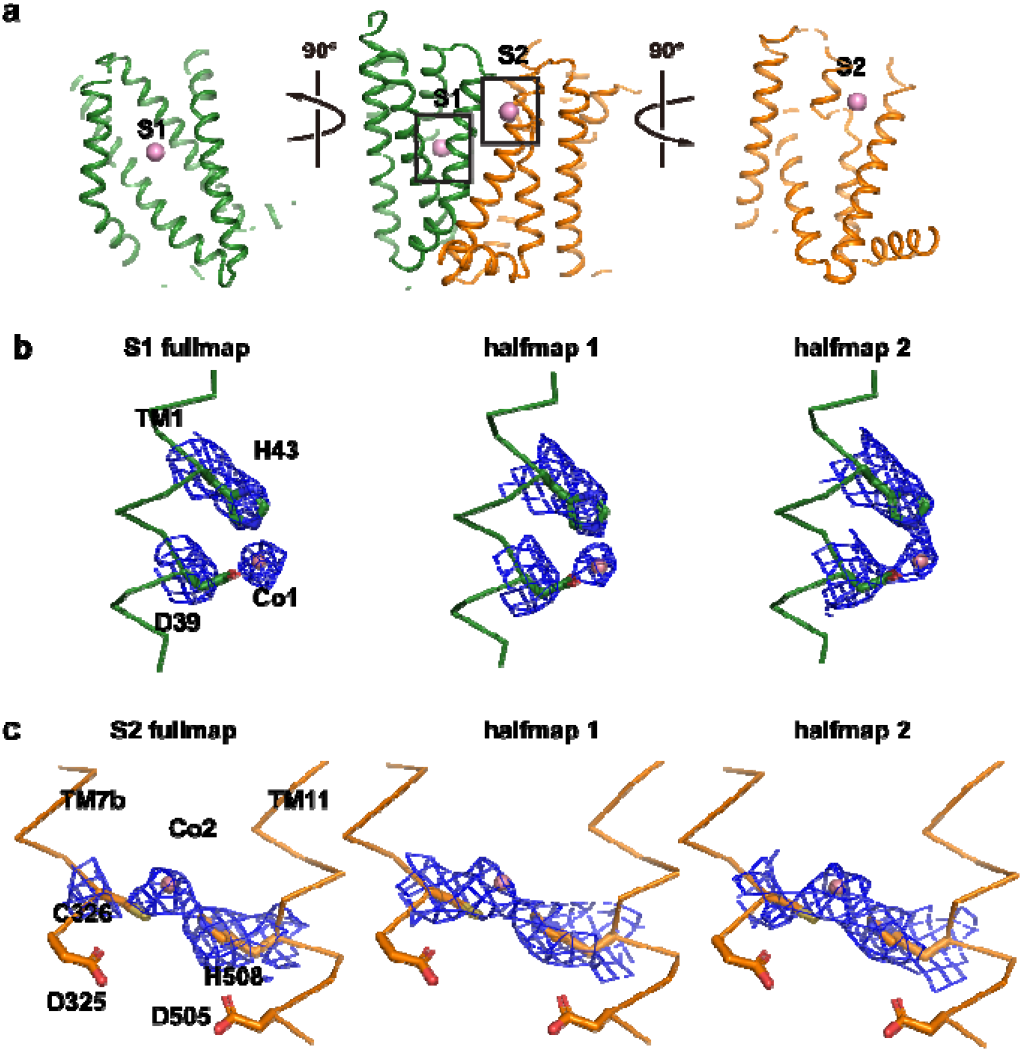
Two ion binding sites in TsFpn. **a.** TsFpn in cartoon representation is shown in three orientations with S1 and S2 marked as sticks. **b.** Density maps of S1 are shown in blue mesh. Part of the TM1 is shown as trace and the side chains of Asp39 and His43 are shown in stick. Co^2+^ is shown as a magenta sphere. **c.** Density maps of S2 are shown in blue mesh. Part of the TM7 and 11 are shown as trace and the side chains of Asp325, Cys326, Asp505 and His508 are shown as sticks.

### TsFpn is an electroneutral H^+^/Fe^2+^ antiporter

As a first test to validate the ion binding sites, we examined pH dependency of metal ion binding and transport in TsFpn because both the S1 and S2 sites contain a histidine. TsFpn does not bind to Co^2+^ in pH 6.0, and the binding affinity gradually recovers as pH increases from 6.0 to 8.0 (Extended Data Figs.1c and 7a-f). Similarly, Co^2+^ uptake is significantly reduced when external pH is 6.5, and the uptake gradually increases as external pH increases from 6.5 to 8.5 (Fig. 4a). These results are consistent with the presence of histidine residues at the ion binding site and provide the first systematic evaluation of pH dependency of ion binding and transport in Fpn.

**Figure 4.**
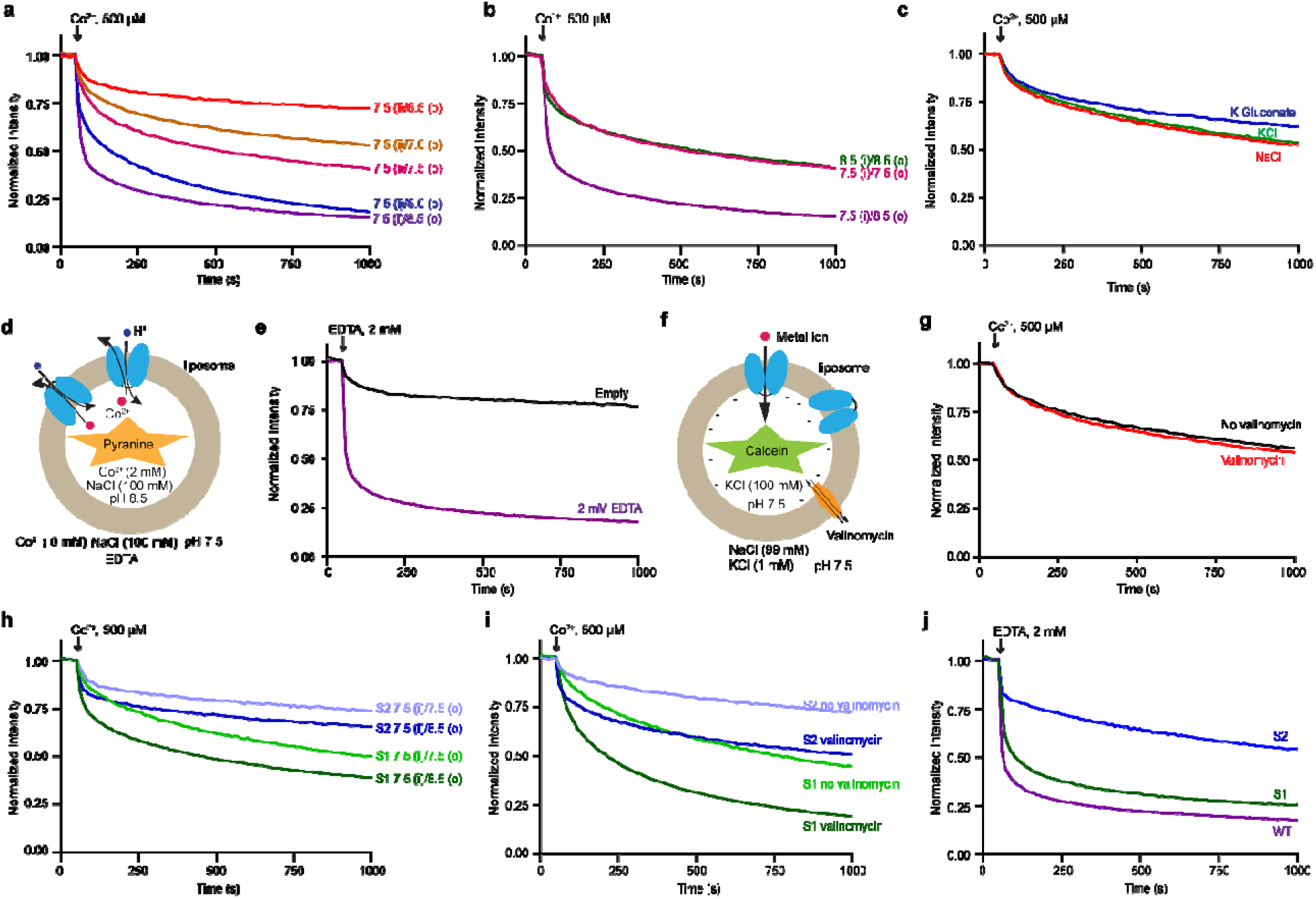
TsFpn is a H^+^/Fe^2+^ antiporter. **a.** Quench of calcein fluorescence over time measured in different external pH from 6.5 to 8.5. The internal pH is 7.5. **b.** Quench of calcein fluorescence over time measured in symmetrical pH, 7.5 inside (i) and 8.5 outside (o). The trace for symmetrical pH 7.5 and the trace for pH 7.5 (internal)/pH8.5 (external) are the same in **a**. **c.** Quench of fluorescence under symmetrical KCl, symmetrical NaCl, and symmetrical K-Gluconate. **d.** Schematic view of a TsFpn containing proteoliposome for monitoring H^+^ influx. **e.** Quench of pyranine fluorescence after addition of EDTA to the external solution. **f.** Schematic view of a TsFpn containing proteoliposome with calcein dye enclosed. KCl is 100 mM inside of the liposomes and 1 mM outside. Valinomycin is added to clamp the membrane potential at ~−120 mV. **g.** Quench of calcein fluorescence over time measured with and without addition of valinomycin under the conditions shown in **f**. **h**. Co^2+^ transport of S1 and S2 mutants under symmetrical and unsymmetrical pH. **i**. Quench of calcein fluorescence over time measured with and without addition of valinomycin under the conditions shown in **f**. **j.** Quench of pyranine fluorescence for S1 and S2 mutants.

Intrigued by the drastically enhanced Co^2+^ uptake at elevated pH (8.0 and 8.5), we wondered whether it was the lower H^+^ concentration or the H^+^ gradient responsible for enhanced metal ion transport. Ion transport under symmetrical pH 8.0 or 8.5 is not significantly different from that in symmetrical pH 7.5 (Fig. 4b), indicating that metal ion uptake is enhanced by the pH gradient, i.e., a higher H^+^ concentration inside of the liposomes. These results suggest that TsFpn is a H^+^/Fe^2+^ antiporter in which metal ion transport is coupled to H^+^ movement in the opposite direction. We further tested this hypothesis in the following three experiments.

We first varied salt composition in the assay buffer and we found that neither Na^+^, K^+^ nor Cl^−^ enhances metal ion transport (Fig. 4c). Second, we measured proton transport directly in a flux assay. In this experiment, uptake of H^+^ is monitored by a proton sensitive fluorescent dye pyranine trapped inside of the liposomes and the H^+^ uptake is driven by efflux of Co^2+^ (Fig. 4d-e and Methods). This result confirms the coupled movement of H^+^ and Co^2+^ in opposite directions.

Third, we determined the stoichiometry of the coupled H^+^ and Co^2+^ movement by examining if metal ion transport in TsFpn is affected by a membrane potential. We set the membrane potential at ~−120 mV by having a 100-fold K^+^ concentration gradient in the presence of a K^+^ selective ionophore valinomycin. Membrane potential of a vesicle is defined assuming 0 mV at outside. We found that the membrane potential has no effect on the rate of either Fe^2+^ or Co^2+^ uptake (Fig. 4f-g). This result indicates that metal ion transport in TsFpn is electroneutral and that the most likely stoichiometry of H^+^ to Fe^2+^ is 2 to 1. Taken together, our results led to the conclusion that TsFpn is an electroneutral 2H^+^/Fe^2+^ antiporter.

### Mutational study of metal ion binding sites

As a second test to validate the ion binding sites, we made mutations to the S1 and S2 binding sites and measured ion binding and transport. Two double mutations were made: the S1 mutation, i.e., Asp39Ala-His43Ala, and the S2 mutation, i.e., Cys326Ala-His508Ala. Both the S1 and S2 mutations can be purified and are stable after purification (Extended Data Fig. 1d-e). We measured metal ion transport in symmetrical pH 7.5 and with a pH gradient (pH7.5 inside and pH 8.5 outside), and we determined if ion transport remains electroneutral.

The S1 mutation has similar transport activity to that of the wild type TsFpn in symmetrical pH 7.5. It has a modest increase in transport activity under the pH gradient conditions and the increase is much smaller than the increase observed in the wild type (Fig. 4b and 4h). This result indicates that the coupled transport of H^+^ and Co^2+^ is affected by the S1 mutation. Consistent with this conclusion, Co^2+^ transport in the S1 mutation becomes electrogenic and shows a large increase in transport activity under a membrane potential of ~−120 mV (Fig. 4i). Enhanced Co^2+^ transport under −120 mV indicates that less than two H^+^ is transported for each Co^2+^, i.e., H^+^ transport is impaired. Since Co^2+^ transport is about similar to that of the wild type in symmetrical pH, it is likely the S1 mutation affects H^+^ transport.

The S2 mutation has significantly lower transport activity than that of the wild type under both the symmetrical and pH gradient conditions. Transport activity is enhanced in the pH gradient, although it is difficult to determine if the enhancement is similar to that of the wild type (Fig. 4h). Co^2+^ transport is also enhanced under −120 mV membrane potential, indicating that less than 2 H^+^ is transported for each Co^2+^ (Fig. 4i). Since Co^2+^ transport is significantly lower in symmetrical pH, it is likely the S2 mutation affects both H^+^ and Co^2+^ transport.

Both S1 and S2 mutations can still transport proton and the effects of the mutations mirror the effects observed in the Co^2+^ transport assay (Fig.4j). Combined, these results led us to conclude that both the S1 and S2 sites are important for H^+^ transport, while for Co^2+^ transport the S2 site is critical and the S1 site seems redundant. Because the S2 mutation maintains a small yet significant level of transport, it is likely that the S1 site could mediate metal ion transport but not as efficiently.

To estimate how the two sites contribute to metal ion binding, we measured Co^2+^ binding to the S1 and S2 mutations by ITC. The S1 mutation binds to Co^2+^ with a K_d_ of 266.3 ± 23.8 μM, and the S2 mutation binds to Co^2+^ with a K_d_ of 162.4 ± 16.0 μM (Extended Data Fig. 7g-h). When both sites are mutated (Asp39Ala/His43Ala/Cys326Ala/His508Ala), the binding affinity becomes 616.0 ± 44.9 μM (Extended Data Figs. 1f and 7i). These results suggest that both S1 and S2 site contribute to metal ion binding.

## Discussion

In summary, we solved the structure of TsFpn in an outward-facing conformation and we identified two potential metal ion binding sites S1 and S2. We also found that TsFpn is a H^+^/Fe^2+^ antiporter and the transport of each Fe^2+^ is coupled to two H^+^. Further studies showed that both S1 and S2 sites are involved in H^+^ transport while the S2 site is more critical to metal ion transport.

We generated a model of TsFpn in the inward facing conformation by aligning the N- and C-domains separately on their equivalent domains in the inward-open bacterial BbFpn structure^21^ (Extended Data Fig. 8a). The transmembrane domains of the two structures align reasonably well with a root mean square distance of 3.6 Å for the N-domain and that 2.1 Å for the C-domain. Both the S1 and S2 sites are solvent accessible in the inward-facing model of TsFpn (Extended Data Fig. 8b), suggesting that a canonical rock-switch type motion of the N- and C-domains could achieve alternating access to the substrates (Fig.5a-d). We speculate that two H^+^ could bind to both S1 and S2 and be transported across the membrane, while one Fe^2+^ could bind to either S1 or S2 sites and be transported in the opposite direction. Further study is required to reveal details of how structural changes during the transport allow the two sites to coordinate and transport H^+^ and Fe^2+^ in opposite directions.

Although TsFpn and BbFpn share the same MFS fold, the S1 and S2 binding sites in TsFpn differ from the ion binding sites identified in BbFpn. The metal ion binding site identified in the initial BbFpn structure is formed by residues Thr20, Asp24, Asn196, Ser199, and Phe200, and these residues correspond to Ser35, Asp39, Asn212, Ser215, and Met216 in TsFpn (Extended Data Figs. 6 and 9a-b). However, only Asp39 is involved in S1 and when the two structures are aligned, the two binding sites are 8.6 Å away (Extended Data Fig. 9a). In a more recent publication, another metal ion binding site was identified in the BbFpn formed by residues Trp254, His261 and Thr386. However, the new metal ion binding site contains an EDTA molecule that helps coordinate a bound metal ion^22^. This site is close to S2 in TsFpn, however, none of the coordinating residues are part of the S2 (Extended Data Figs. 6 and 9c). Bacterial BbFpn was also shown to have higher rate of uptake when external pH increases, but the enhanced transport was due to the pH and not the pH gradient^21^. This result indicates that BbFpn has a different metal ion transport mechanism.

We mapped known missense mutations that cause ferroportin diseases onto the TsFpn structure (Fig. 5e). The structure provides insights into how certain mutations may lead to diseases. For example, six mutations are mapped to regions close to S1 or S2 and these mutations likely affect ion binding. Eighteen mutations are mapped to regions where the N- and C-domain make contact in the current structure and these mutations likely affect transport activity. Moreover, we show that transport activity of TsFpn is inhibited by human hepcidin, and we mapped residues known to affect hepcidin inhibition to the TsFpn structure^3^. These residues cluster mostly on the extracellular surface with a few deeper into the cavity including Cys326 of the S2 site. It appears that hepcidin could perturb either H^+^ or Fe^2+^ binding by interacting with the S2 site.

**Figure 5.**
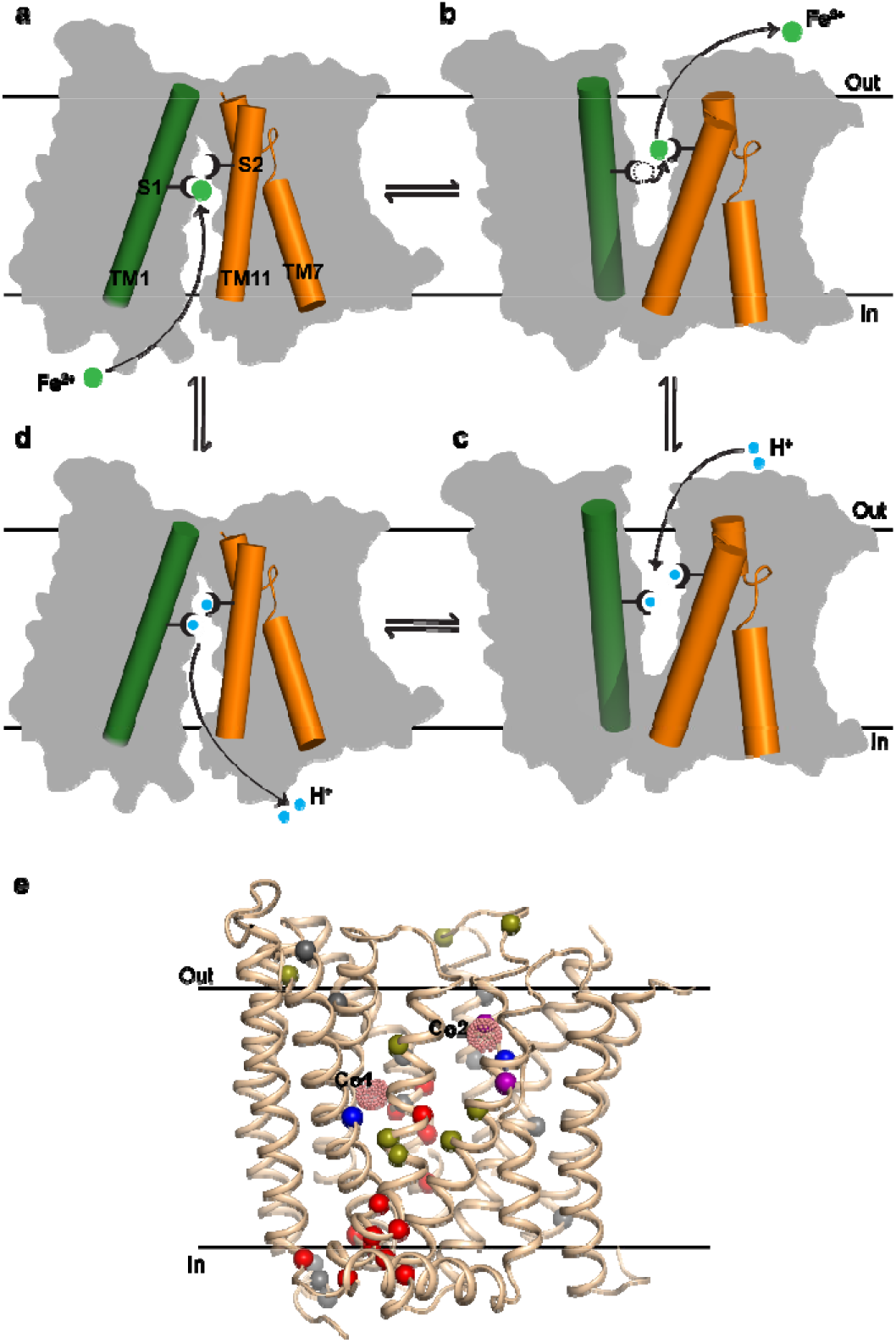
Proposed mechanism of ion transport and disease-related mutations in Fpn. **a-d.** Proposed conformational changes of Fpn during the transport cycle. A cytosolic Fe^2+^ ion (green sphere) binds to either the S1 or S2 sites (drawn as forks) of a Fpn at the inward-facing conformation (**a**); Fpn then switches to the outward-facing conformation to allow the bound Fe^2+^ to escape to the extracellular side (**b**). Two protons bind to the S1 and S2 sites in an outward-facing conformation of Fpn (**c**) and enter the cytosol when Fpn returns to the inward-facing conformation (**d**). **e.** Cartoon representation of TsFpn. The bound Co^2+^ ions are shown as pink dots. Cα of disease-related missense mutations are shown as spheres. Mutations close to the ion binding sites are colored blue, those at the interface between the N and C domains are colored red, and ones that are reported to affect hepcidin binding and endocytosis are colored olive. Cys326 and His508 are shown in purple. All other mutations are colored in grey.

## Methods

### Cloning, expression, and purification of TsFpn

The Fpn gene (accession number XP_008060857) from *Carlito syrichta* (*Tarsius syrichta*, Philippine tarsier) was codon-optimized and cloned into a pFastBac dual vector^32^ for production of baculovirus by the Bac-to-Bac method (Invitrogen). High Five Cells (Thermofisher) at a density of ~3×10^6^ cells/ml were infected with baculovirus and grown at 27 °C for 60-70 hour before harvesting. Cell membranes were prepared following a previous protocol^32^ and frozen in liquid nitrogen.

Purified membranes were thawed and homogenized in 20 mM HEPES, pH 7.5, 150 mM NaCl and 2mM β-mercaptoethanol, and then solubilized with 1% (w/v) Lauryl Maltose Neopentyl Glycol (LMNG, Anatrace) at 4 °C for 2 h. After centrifugation (55,000g, 45min, 4 °C), TsFpn was purified from the supernatant using a cobalt-based affinity resin (Talon, Clontech) and the His_6_-tag was cleaved by TEV protease at room temperature for 1 hour. TsFpn was then concentrated to 3-6 mg/ml (Amicon 50 kDa cutoff, Millipore) and loaded onto a size-exclusion column (SRT-3C SEC-300, Sepax Technologies, Inc.) equilibrated with 20 mM HEPES, pH7.5, 150 mM NaCl, 1 mM (w/v) n-dodecyl-β-D-maltoside (DDM, Anatrace).

Mutations to TsFpn were generated using the QuikChange method (Stratagene) and the entire cDNA was sequenced to verify the mutation. Mutants were expressed and purified following the same protocol as wild type.

### Generation of monoclonal antibodies and Fab fragments

Monoclonal antibodies against the TsFpn (IgG2b, κ) were raised using standard methods (Monoclonal Core, Vaccine and Gene Therapy Institute, Oregon Health & Science University). High affinity and specificity of the antibodies for properly folded TsFpn was assayed by ELISA and western blot (no binding). Three out of twenty antibodies were selected for large scale production. Fab fragments were generated by papain cleavage of whole antibody at a final concentration of 1 mg/mL for 2 hours at 37 °C in 50 mM Phosphate buffer saline, pH 7.0, 1 mM EDTA, 10 mM cysteine and 1:50 w:w papain:antibody. Digestion was quenched using 30 mM iodoacetamide at 25 °C for 10 min. Fab was purified by anion exchange using a Q Sepharose (GE Healthcare) column in 10 mM Tris, pH 8.0 and a NaCl gradient elution. TsFpn-Fab complexes were further verified by size-exclusion chromatography (shift in elution volume and SDS-PAGE) and 11F9 was selected for structural studies.

### TsFpn-11f9(Fab) complex

Purified TsFpn was mixed with the 11F9 Fab at 1:1.1 molar ratio and incubated 30 min on ice. The complex was then concentrated to 3-6 mg/ml (Amicon 100 kDa cutoff, Millipore) and loaded onto a size-exclusion column equilibrated with 20 mM HEPES, pH7.5, 150 mM NaCl, 1 mM n-dodecyl-β-D-maltoside (DDM, Anatrace). The TsFpn-Fab complex was used in the ITC measurement of Co^2+^ binding and in nanodisc reconstitution for cryo-EM grid preparations.

### Nanodisc reconstitution

MSP1D1 was expressed and purified following an established protocol^33^. For lipid preparation, 1-palmitoyl-2-oleoyl-sn-glycero-3-phospho-(1’-rac)-choline (POPC, Avanti Polar Lipids), 1-palmitoyl-2-oleoyl-sn-glycero-3-phospho-(1’-rac)-ethanolamine (POPE, Avanti Polar Lipids) and 1-palmitoyl-2-oleoyl-sn-glycero-3-phospho-(1’-rac)-glycerol (POPG, Avanti Polar Lipids) were mixed at a molar ratio of 3:1:1, dried under Argon and resuspended with 14 mM DDM^34^. For nanodisc reconstitution, TsFpn, Fab of 11f9, MSP1D1 and lipid mixture were mixed at a molar ratio of 1:(1.1):(2.5):(62.5) and incubated on ice for 1 hour. Detergents were removed by incubation with Biobeads SM2 (Bio-Rad) overnight at 4 °C. The protein lipid mixture was loaded onto a size-exclusion column equilibrated with 20 mM HEPES, pH7.5, 150 mM NaCl. The purified nanodisc elutes at 13.6 ml and was concentrated to 13 mg/ml and incubated with 10 mM CoCl_2_ for 30 min before cryo-EM grid preparation.

### Cryo-EM sample preparation and data collection

The cryo grids were prepared using Thermo Fisher Vitrobot Mark IV. The Quantifoil R1.2/1.3 Cu grids were glow-discharged with air for 15 sec at 10 mM in a Plasma Cleaner (PELCO EasiGlow™). Aliquots of 3.5 μl purified TsFpn-11f9 in nanodisc were applied to glow-discharged grids. After being blotted with filter paper (Ted Pella, Inc.) for 4.0 s, the grids were plunged into liquid ethane cooled with liquid nitrogen. A total of 1838 micrograph stacks were collected with SerialEM^35^ on a Titan Krios at 300 kV equipped with a K2 Summit direct electron detector (Gatan), a Quantum energy filter (Gatan) and a Cs corrector (Thermo Fisher), at a nominal magnification of 105,000 × and defocus values from −2.0 μm to −1.2 μm. Each stack was exposed in the super-resolution mode for 5.6 s with an exposing time of 0.175 s per frame, resulting in 32 frames per stack. The total dose rate was about 50 e^−^/Å^2^ for each stack. The stacks were motion corrected with MotionCor2^36^ and binned 2 fold, resulting in a pixel size of 1.114 Å/pixel. In the meantime, dose weighting was performed^37^. The defocus values were estimated with Gctf^38^.

### Cryo-EM data processing

A total of 1,246,999 particles were automatically picked with RELION 2.1^39–41^. After 2D classification, a total of 946,473 particles were selected and subject to a global angular search 3D classification with one class and 40 iterations. The outputs of the 35th-40th iterations were subjected to local angular search 3D classification with four classes separately. A total of 571,511 particles were selected by combining the good classes of the local angular search 3D classification. After handedness correction, a skip-align classification procedure was performed to further classify good particles, yielding a total of 215,752 particles, which resulted into a reconstruction with an overall resolution of 3.1 Å after 3D auto-refinement with an adapted mask. The resolution of the map was further improved to 3.0 Å after Bayesian polishing^42^.

All 2D classification, 3D classification, and 3D auto-refinement were performed with RELION 2.1 or RELION 3.0. Resolutions were estimated with the gold-standard Fourier shell correlation 0.143 criterion^43^ with high-resolution noise substitution^44^.

### Model building and refinement

For *de novo* model building of TsFpn-11F9 complex, a ploy-Alanine model was first manually built into the 3.0 Å density map in COOT^45^ and side chains were added next. Structure refinements were carried out by PHENIX in real space with secondary structure and geometry restraints^46^. The EMRinger Score was calculated as described^47^.

### Proteoliposome preparation

POPE and POPG (Avanti Polar Lipids) was mixed at 3:1 molar ratio, dried under Argon and vacuumed overnight to remove chloroform. The lipid was resuspended in the reconstitution buffer (20 mM HEPES, pH 7.5, 100 mM NaCl) to a final concentration of 10 mg/ml, sonicated to transparency and incubated with 40 mM n-decyl-β-D-maltoside (DM, Anatrace) for 2 h at room temperature under gentle agitation. Wild type or mutant TsFpn was added at 1:100 (w/w, protein:lipid) ratio. The detergent was removed by dialysis at 4 °C against the reconstitution buffer. Dialysis buffer was changed once a day and the liposomes were harvested after 4 days, aliquoted, and frozen at −80 °C.

### Divalent metal ion flux assay

Liposomes were mixed with 250 μM calcein and frozen-thawed three times. After the liposomes were extruded to homogeneity with 400 nm filter (NanoSizer™ Extruder, T&T Scientific Corporation), free calcein was removed through a desalting column (PD-10, GE Healthcare) equilibrated with the dialysis buffer. Calcein fluorescence was monitored in a quartz cuvette at 37°C. Fluorescence was monitored in a FluoroMax-4 spectrofluorometer (HORIBA) with 494 nm excitation and 513 nm emission at 10 s internals. The transport was initiated by the addition of 0.5 mM CoCl_2_ or 100 μM fresh prepared FeSO_4_. When Fe^2+^ is used, 1 mM vitamin C was added in the external solution. The rate of ion transport is estimated by the initial slope of the traces of fluorescence quench.

In experiments when internal solution needs to be replaced, liposomes were centrifuged at 47000g for 30 min and resuspended in a desired internal solution. A fluorescent dye was then loaded into the liposomes by the same freeze-thaw processes and free dye was removed by a desalting column.

### Pyranine assay

Liposomes were centrifuged at 47000g for 30 min and resuspended in inside buffer (5 mM Tris, pH 8.5, 100 mM NaCl). Liposomes were mixed with 250 μM pyranine and 2 mM CoCl_2_ and frozen-thawed three times. After the liposomes were extruded to homogeneity with 400 nm filter (NanoSizer™ Extruder, T&T Scientific Corporation), free dye was removed through a desalting column (PD-10, GE Healthcare) equilibrated with the outside buffer (5 mM HEPES, pH 7.5, 100 mM NaCl, 2 mM CoCl_2_). Pyranine fluorescence was monitored in a quartz cuvette at 37°C in a FluoroMax-4 spectrofluorometer (HORIBA) with 460 nm excitation and 510 nm emission at 10 s internals. The transport was initiated by the addition of 2 mM EDTA.

### Isothermal titration calorimetry

Protein samples were purified as described above and concentrated to around 50-100 μM (3-6 mg/ml). TsFpn was in the ITC buffer that contains 20 mM HEPES pH 7.5, 150 mM NaCl, 1 mM DDM. The ITC measurements were performed with a Nano ITC microcalorimeter (TA Instruments) at 25 °C. CoCl_2_ stock at 5 mM was prepared in the same ITC buffer injected 25 times (1.01 μl for injection 1 and 2.02 μl for injections 2–15), with 175 s intervals between injections. The background data obtained from injecting Co^2+^ into the ITC buffer were subtracted before the data analysis. The data were fitted using the Origin8 software package (MicroCal). Measurements were repeated three times.

## Acknowledgments

This work was supported by grants from NIH (DK122784, HL086392 and GM098878 to MZ), Cancer Prevention and Research Institute of Texas (R1223 to MZ). Ara Parseghian Medical Research Foundation (to N.Y.). N.Y. is supported by the Shirley M. Tilghman endowed professorship from Princeton University. We thank Paul Shao for technical support during EM image acquisition. We acknowledge the use of Princeton’s Imaging and Analysis Center, which is partially supported by the Princeton Center for Complex Materials, and the National Science Foundation (NSF)-MRSEC program (DMR-1420541).

## Data Availability

The atomic coordinates of TsFpn-Fab complex have been deposited in the PDB (http://www.rcsb.org) under the accession codes 6VYH. The corresponding electron microscopy maps have been deposited in the Electron Microscopy Data Bank (https://www.ebi.ac.uk/pdbe/emdb/) under the accession codes EMD-21460.

## Author Contributions

M.Z., Z.R., Y.P., and J.S. conceived the project. S.G. led the effort of cryo-EM grid preparation, data collection and analysis and was assisted by Z.R., J.S. and L.W.. Y.P., Z.R., J.S., L.W., Y.Y., H.Z., Z.X., P.B. and A.L. conducted experiments. Y.P., Z.R., G.S., J.S., Z.X., P.B., A.L., N.Y. and M.Z. analyzed data. Z.R., J.S., Y.P. and M.Z. wrote the initial draft and all authors participated in revising the manuscript.

## Extended Data

**Extended Data Figure 1.**
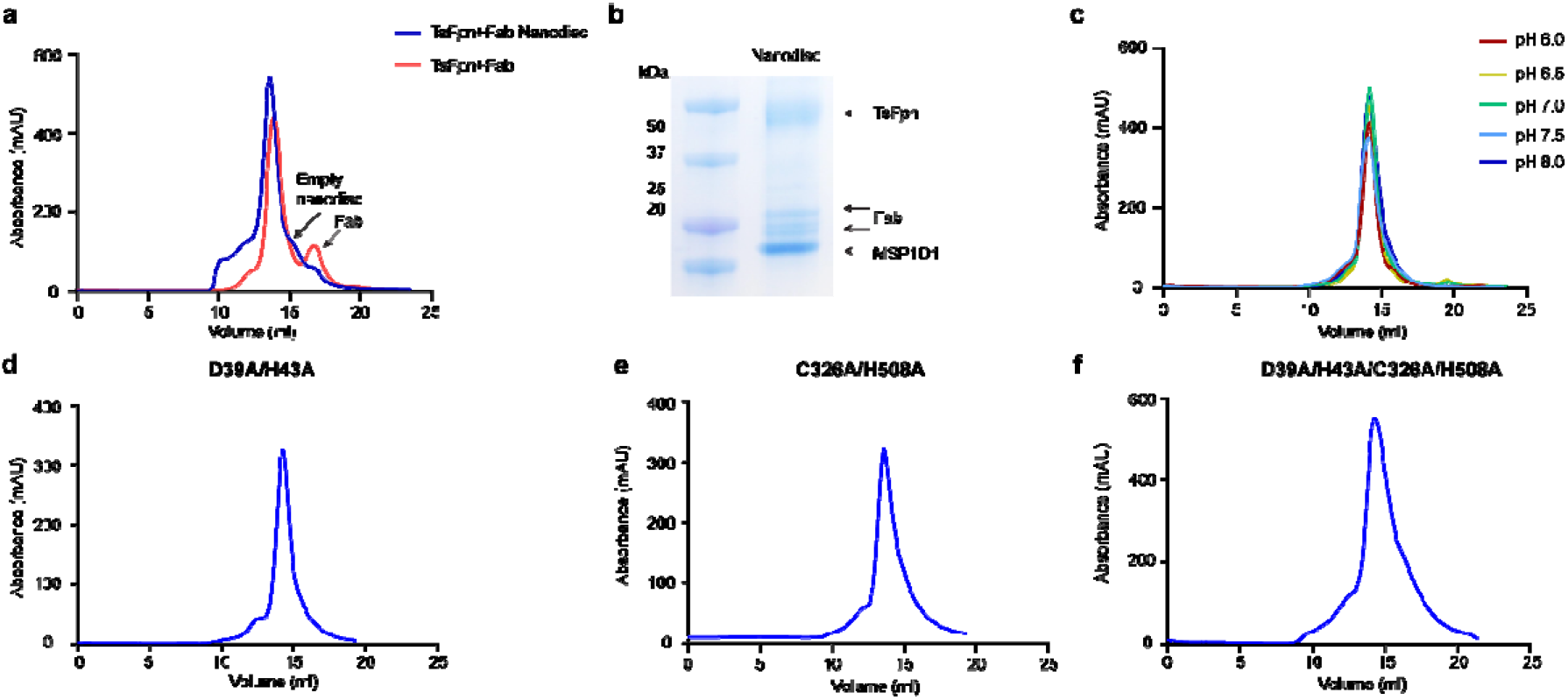
Wild type and mutant TsFpn proteins. **a.** Size-exclusion chromatography of TsFpn in complex with Fab of 11F9 before and after reconstitution into nanodiscs. **b.** SDS-PAGE of the reconstitution. **c.** Size-exclusion chromatography of TsFpn in pH ranging from 6.0 to 8.5. **d-f.** Size-exclusion chromatography of TsFpn with S1 (D39A/H43A), S2 (C326A/H508A) and S1+S2 (D39A/H43A/C326A/H508A) mutations.

**Extended Data Figure 2.**
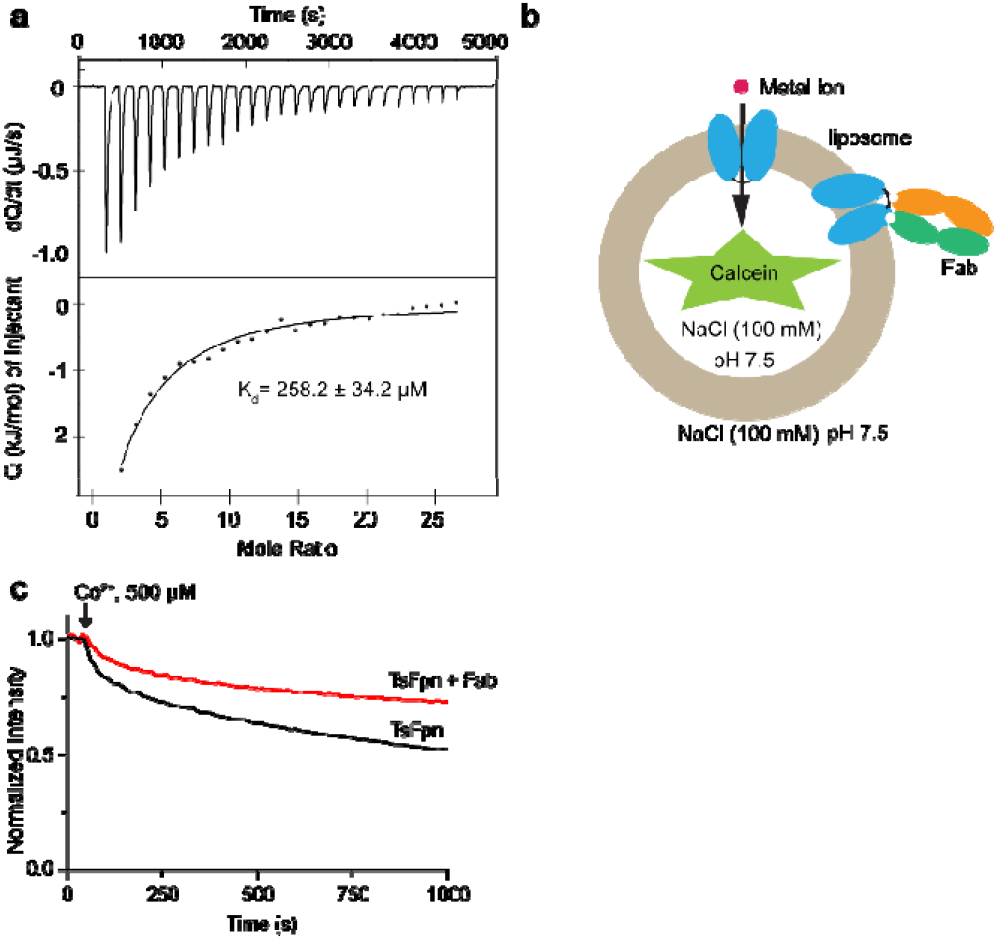
Effect of Fab on Co^2+^ binding and transport. **a.** ITC measurement of Co^2+^ binding to TsFpn in the presence of 11F9 Fab. **b.** Schematic illustration of a proteoliposome with TsFpn in both orientations. The liposomes are loaded with calcein and the 1 μM of 11F9 Fab was added to the external side of the liposomes. **c.** Quench of calcein fluorescence in the absence (black trace) and presence of 1 μM 11F9 Fab (red trace).

**Extended Data Figure 3.**
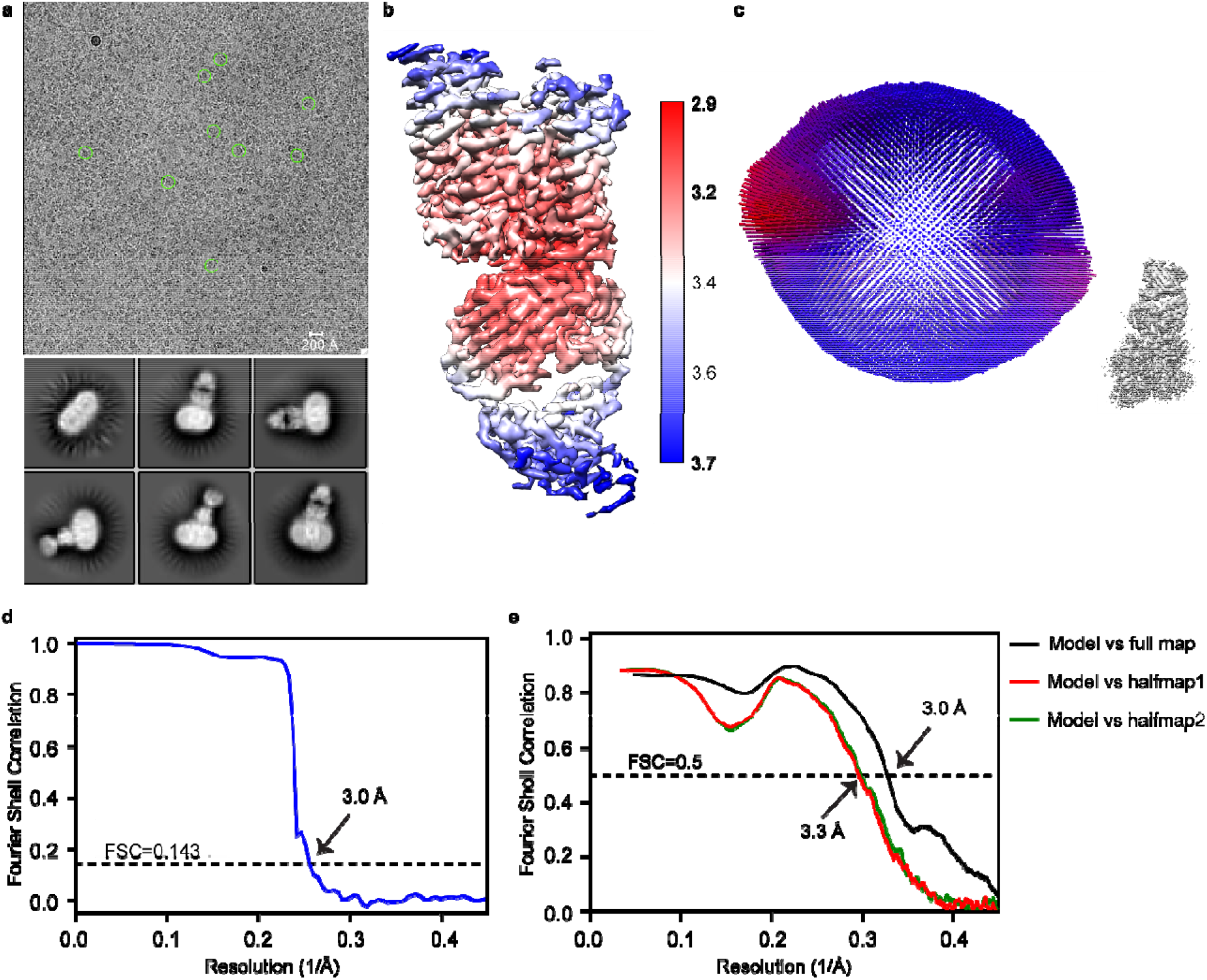
Cryo-EM analysis of the TsFpn-Fab complex reconstituted in nanodisc. **a.** Representative electron micrograph and 2D class averages of cryo-EM particle images. **b.** Local resolution map for the 3D reconstruction of the TsFpn-Fab complex. **c.** Euler angle distribution of the TsFpn-Fab complex in the final 3D reconstruction. **d.** The gold-standard Fourier shell correlation curve for the final map. **e.** FSC curve of the refined model of the TsFpn-Fab complex versus the full map (black) and individual halfmaps (red and green).

**Extended Data Figure 4.**
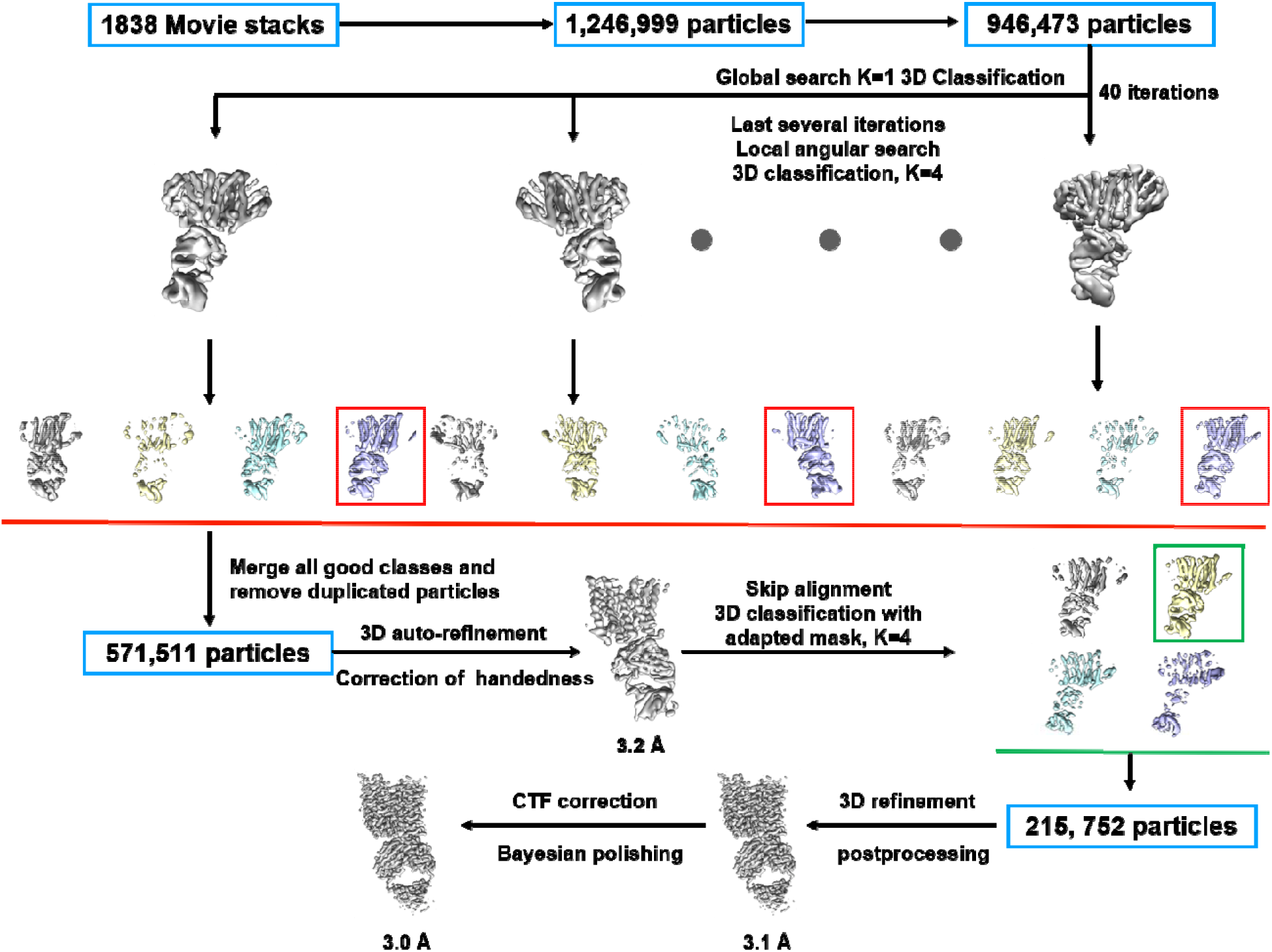
Flow chart of Cryo-EM data processing.

**Extended Data Figure 5.**
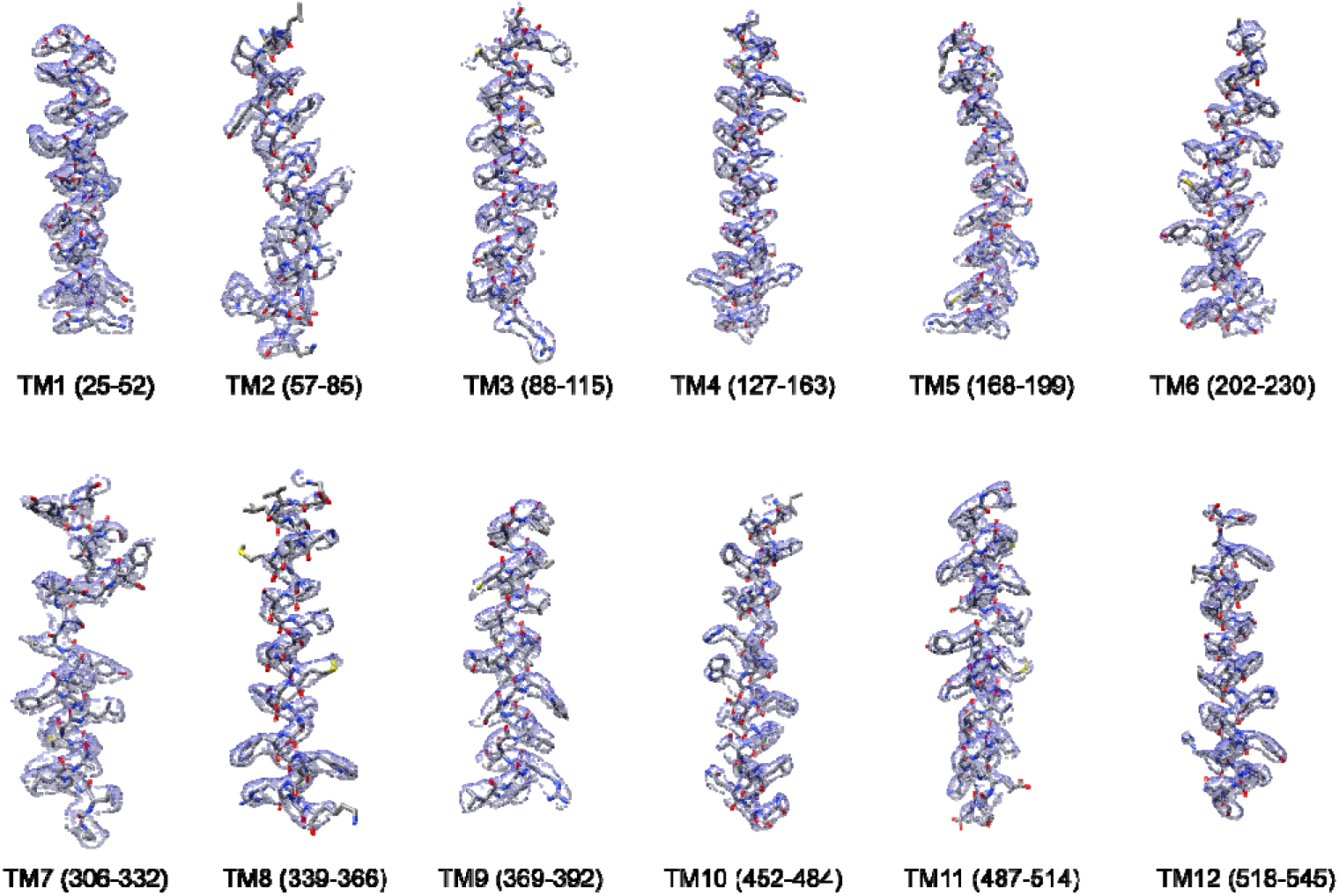
Representative densities of TsFpn.

**Extended Data Figure 6.**
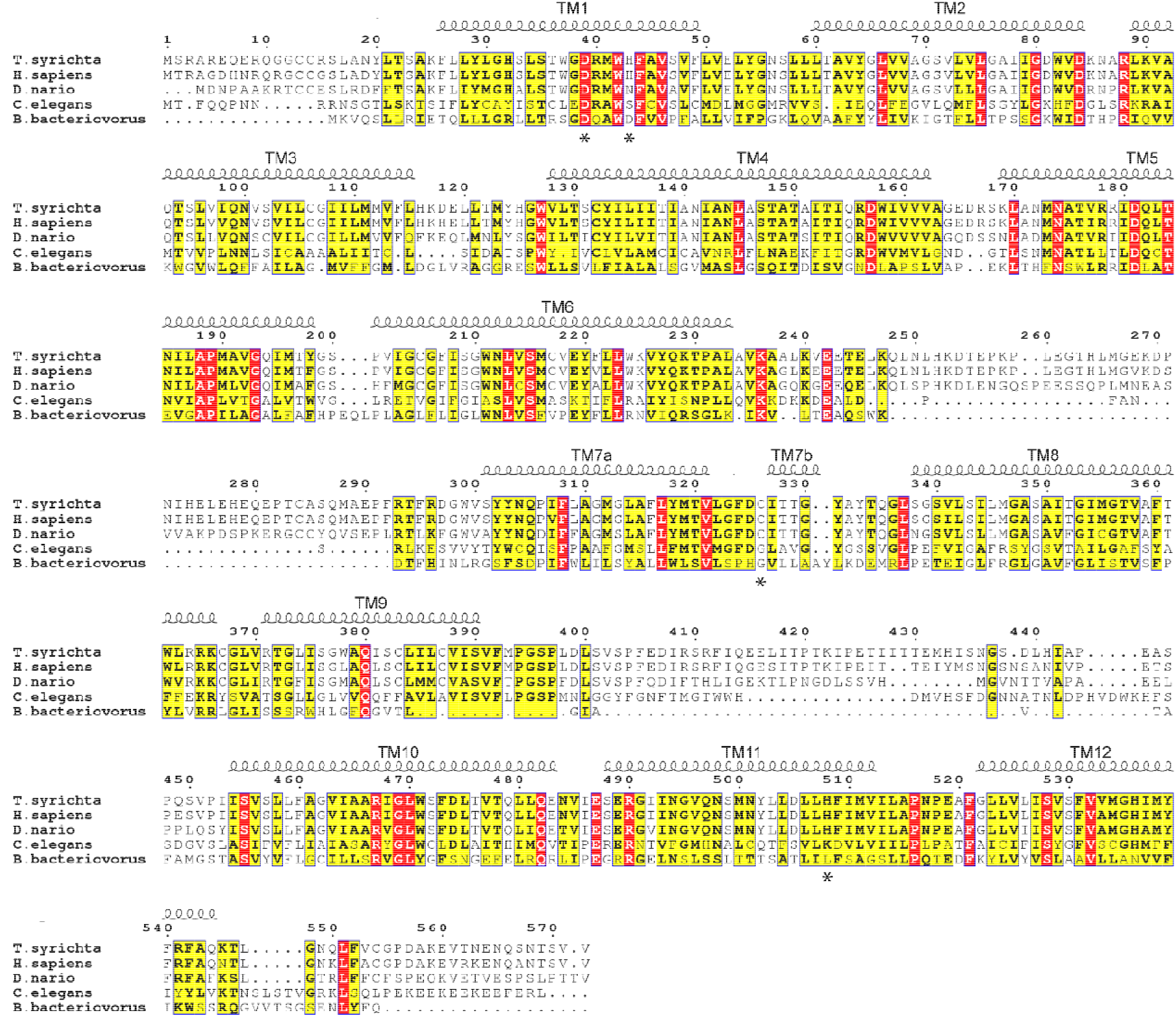
Fpn sequence alignment. **a.** Fpn from Philippine tarsier (Uniprot accession number A0A1U7U6F1), human (Q9NP59), zebrafish (Q9I9R3), worm (Q8IA95), and BbFpn (Q6MLJ0) are aligned using the Clustal Omega server^50^. Secondary structural elements of Fpn are marked above the alignment. Residues are colored based on their conservation using the ESPript server^51^. Residues at the two metal binding sites are labeled with stars.

**Extended Data Figure 7.**
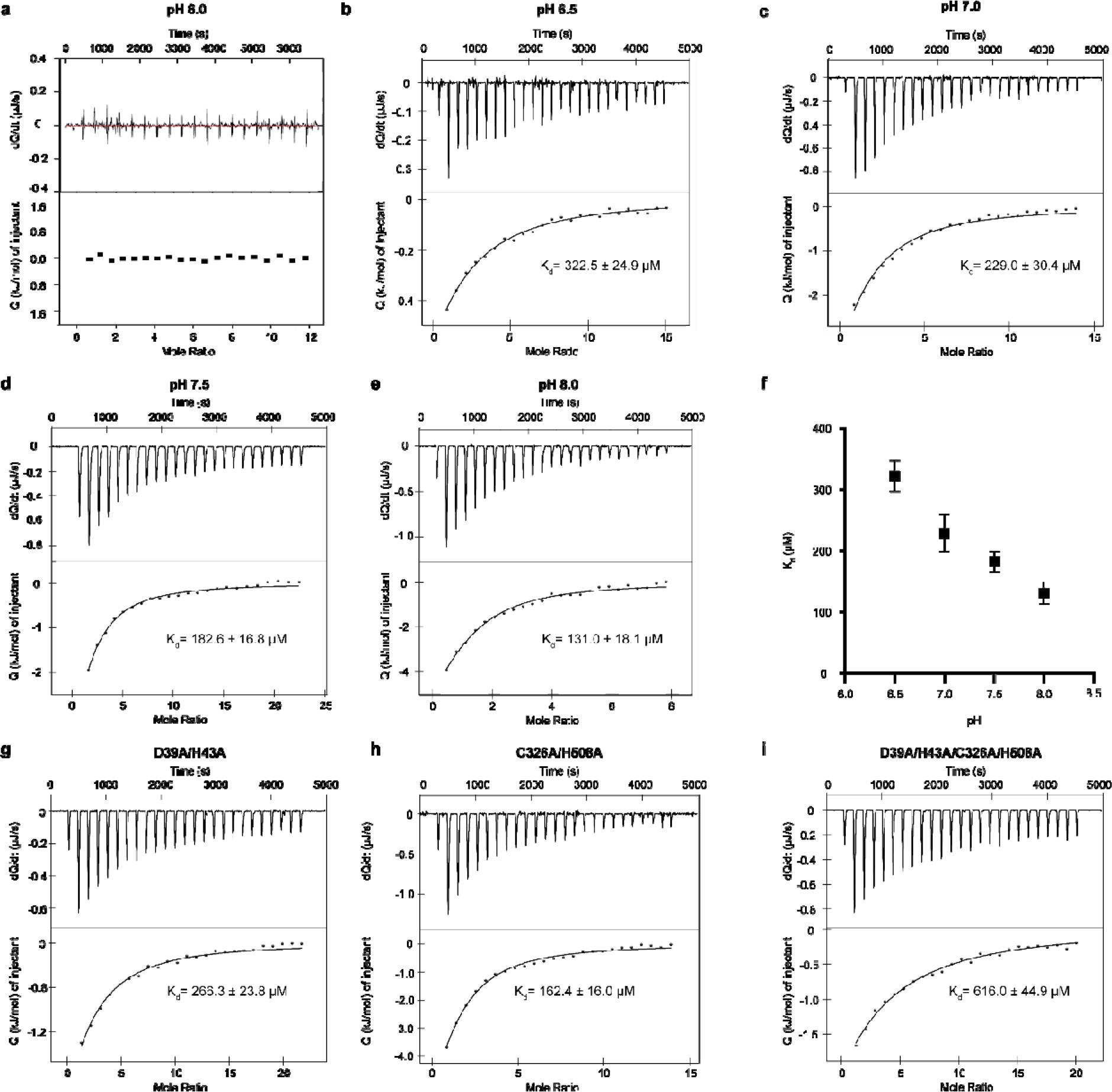
ITC measurements of Co^2+^ binding in different pH. ITC measurements of Co^2+^ binding to TsFpn in pH 6.0 (**a**); pH 6.5 (**b**); pH 7.0 (**c**); pH 7.5 (**d**); pH 8.0 (**e**). **f.** K_d_ of Co^2+^ versus different pH. Error bars are s.e.m.. In **a-e**, the top panel is the rate of heat release versus time and the bottom panel is heat from each injection versus the molar ratios between Co^2+^ and TsFpn. Data points in **b-e** were fit with a single-binding site equation. ITC binding experiment at each pH was repeated three times the composite K_d_ values were plotted in **f. g-i.** ITC measurements of Co^2+^ binding to S1**(g)**, S2 **(h)** and S1+S2 **(i)** mutants.

**Extended Data Figure 8.**
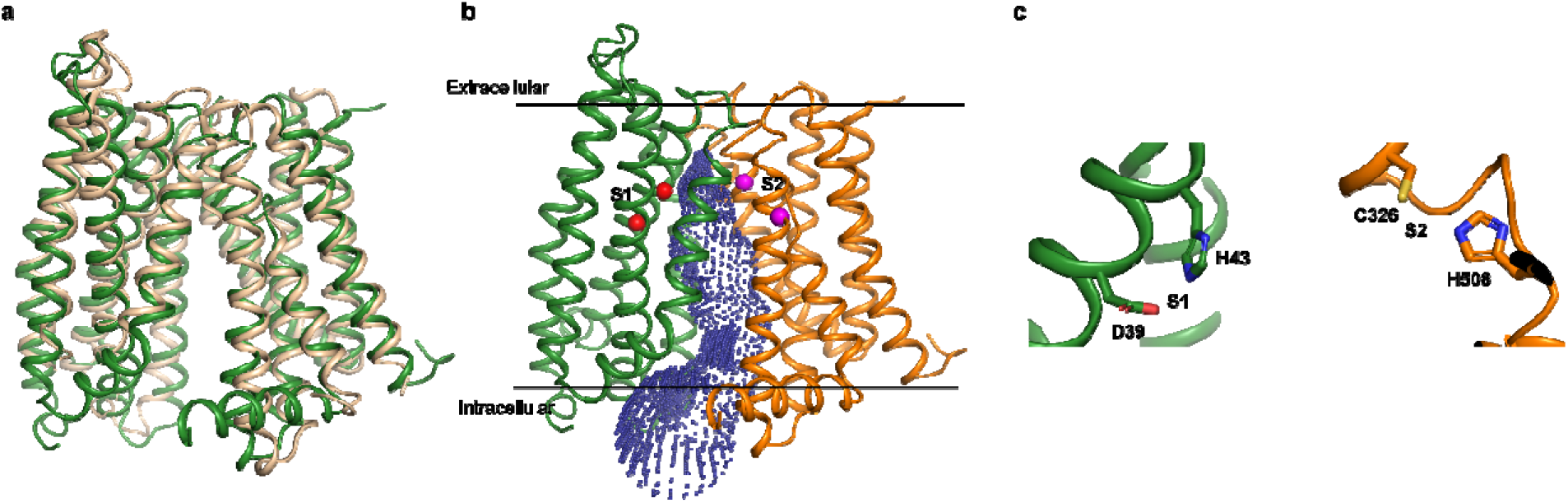
Inward-facing model of TsFpn. **a.** The inward-facing model of TsFpn (cartoon in green) is superposed onto BbFpn (cartoon in wheat) in two views. **b.** cartoon representation of the inward-facing model of TsFpn with the N- and C-domains shown in green and orange, respectively. The solvent accessible regions in the cavity, calculated by HOLE^52^, is shown as blue dots. C-alphas of S1 and S2 are shown as spheres and colored red and magenta, respectively. **c.** Close view of S1 and S2 with side chains shown as sticks.

**Extended Data Figure 9.**
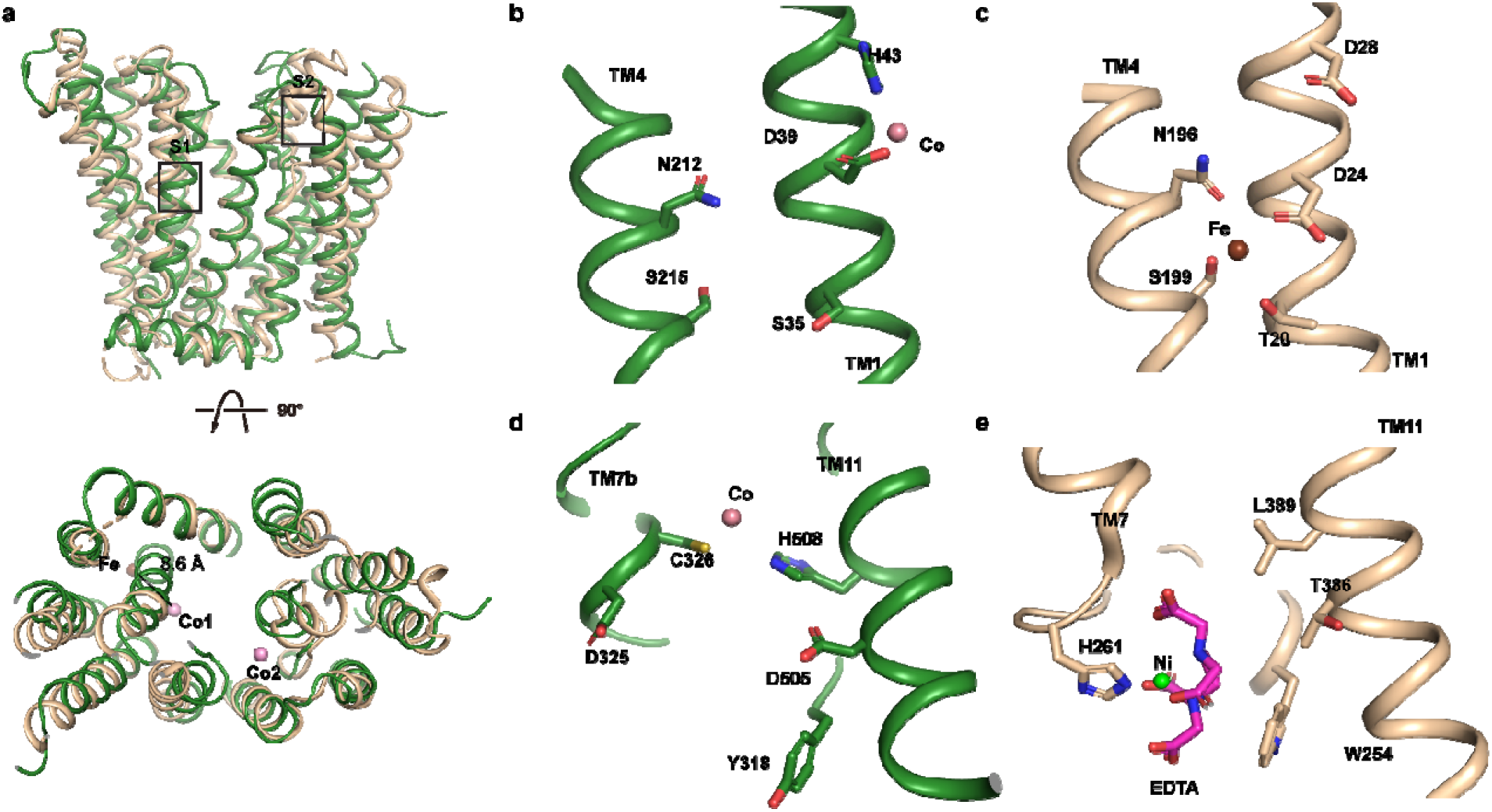
Comparison of ion binding sites in TsFpn and BbFpn. **a.** TsFpn (cartoon in green) is superposed onto BbFpn (cartoon in wheat) in two views. **b and d.** Ion binding sites in TsFpn. **c and e.** Ion binding sites in BbFpn.

**Extended Data Table 1.** Summary of Cryo-EM data collection, processing and structure refinement.

| Protein | TsFpn-Fab |
| --- | --- |
| **Cryo-EM Data Collection** |  |
| Voltage (kV) | 300 |
| Magnification (x) | 105,000 |
| Pixel Size (Å) | 1.114 |
| Electron exposure (e-/Å<sup>2</sup>/frame) | 1.56 |
| Defocus range (µm) | [-2.0, -1.2] |
| Number of image stacks | 1838 |
| Number of frames per stack | 32 |
| **Cryo-EM Data Processing** |  |
| Initial number of particles | 946,473 |
| Final number of particles | 215,752 |
| Symmetry imposed | C1 |
| Map sharpening B factor (Å<sup>2</sup>) | -100 |
| Map resolution (Å) | 3.0 |
| Map resolution range (Å) | 2.9-3.7 |
| FSC threshold | 0.143 |
| **Model Refinement** |  |
| Number of amino acids | 847 |
| Total non-hydrogen atoms | 5392 |
| Average B factor (Å<sup>2</sup>) | 49.83 |
| Bond length r.m.s.d. (Å) | 0.005 |
| Bond angle r.m.s.d. (°) | 0.814 |
| Ramachandran Plot |  |
| Favored (%) | 92.65 |
| Allowed (%) | 7.35 |
| Outliers (%) | 0.00 |
| Rotamer outliers (%) | 0.28 |
| EMRinger Score | 2.56 |

